## Supplementary results, figures, and tables for "Comparative neuroimaging of sex differences in human and mouse brain anatomy"

**Supplementary Materials**

1. **Methods**

**1.1 Demographics information for humans and mice**

Demographics information for the human (***Supplementary Table 1***) and mouse (***Supplementary Table 2***) samples used in this study can be found in the tables below. In ***Supplementary Table 3*** we outline the origin lab for the mouse cohorts used in this study.

***Supplementary Table 1.*** ***Demographics for human sample***

|  |  | **Females** | **Males** | **Statistics** |
| --- | --- | --- | --- | --- |
| **Sample size** | N | 516 | 454 |  |
| **Age** | *Mean* | 29.41 | 27.9 | F(1,1082)=57.13, p=8.67e-14 *** |
|  | *SD* | 3.68 | 3.61 |  |
|  | *Range* | 22-36 | 22-37 |  |
| **Education (in years)** | *Mean* | 14.96 | 14.84 | F(1,1082)=1.465,p=0.226 |
|  | *SD* | 1.83 | 1.77 |  |
|  | *Range* | 11 to 17 | 11 to 17 |  |
| **Euler number** | *Mean* | -52.47 | -58.26 | F(1,1082)=25.72, p=4.65e-07 *** |
|  | *SD* | 17.66 | 19.25 |  |
|  | *Range* | -126 to -16 | -136 to -16 |  |
| **Zygosity** | *Monozygotic* | 102 | 49 | X^2^= 26.281, p= 8.33e-06 |
|  | *Dizygotic* | 163 | 131 |  |
|  | *Not Twin* | 251 | 274 |  |

**P* < 0.01 **P < 0.001 for ANOVA test of significant difference between groups (males vs. females). SD=standard deviation.

***Supplementary Table 2.*** ***Demographics for mouse sample*.** *For details regarding origin of each mouse cohort refer to* ***Supplementary Table 1.***

|  | **Female** | **Male** | **Statistics** |
| --- | --- | --- | --- |
| **Sample size** | 213 | 216 |  |
| **Age** | | | |
| *Mean* | 62.0 | 62.8 | F(1,70)=0.78, p=0.38 |
| *SD* | 7.5 | 8.6 |  |
| *Range* | 56-90 | 56-90 |  |
| **Background Strain** | | | |
| *C57BL-6J* | 134 | 141 |  |
| *C57BL-6N* | 79 | 75 |  |
| **Mouse Cohort for C57BL6J** | | | |
| A | 10 | 12 | X^2^=5.46, p= 0.91 |
| B | 15 | 15 |  |
| C | 27 | 29 |  |
| D | 13 | 7 |  |
| E | 8 | 11 |  |
| F | 9 | 10 |  |
| G | 7 | 9 |  |
| H | 10 | 10 |  |
| I | 7 | 9 |  |
| J | 10 | 6 |  |
| K | 9 | 8 |  |
| L | 9 | 15 |  |
| **Mouse Cohort for C57BL6N** | | | |
| M | 13 | 19 | X^2^=5.745 p= 0.332 |
| N | 10 | 8 |  |
| O | 25 | 13 |  |
| P | 9 | 9 |  |
| Q | 9 | 13 |  |
| R | 13 | 12 |  |

***Supplementary Table 3****. Information about origin laboratory of WT mice included in the study.*

| **Background Strain** | **Study Cohort Key** | **Origin Laboratory** |
| --- | --- | --- |
| **C57BL6J** | A | University of Michigan; Dr. Diane Robinson |
|  | B | KAIST; Dr. Eunjoon Kim |
|  | C | University of Western Ontario; Dr. Nathalie Berube |
|  | D | UT Southwestern; Dr. Genevieve Konopka |
|  | E | Duke University; Dr. Christelle Golzio |
|  | F | Duke University; Dr. Christelle Golzio |
|  | G | Lost Angeles Children’s Hospital; Dr. Pat Levitt |
|  | H | Columbia University; Dr. Jeremy Veenstra-VenderWeele |
|  | I | Scripps Research Institute; Dr. Gavin Rumbaugh |
|  | J | McMaster University; Dr. Karun Singh |
|  | K | McMaster University; Dr. Jane Foster |
|  | L | University of Toronto Center for Phenogenomics |
| **C57BL6N** | M | The Hospital for Sick Children; Dr. Lauryl Nutter |
|  | N | UC Davis - MIND Institute; Dr. Alex Nord |
|  | O | The Hospital for Sick Children; Dr. Lauryl Nutter |
|  | P | UCSD; Dr. Lilia Iakoucheva |
|  | Q | UT Southwestern; Dr. Graig Powell |
|  | R | UC Davis; Dr. Alexander Nord |

**1.2 Deformation based morphometry for hypothalamus segmentation**

T1-weighted structural MRI scans were converted to the MINC file format (<https://www.mcgill.ca/bic/software/minc>) and processed using the minc-bpipe-library preprocessing pipeline (<http://github.com/CobraLab/minc-bpipe-library>) which performs an iterative whole-brain bias field correction using N4ITK (Avants et al. 2011). Images were cropped to remove excess data around the head to improve subsequent image processing steps. A brain mask was computed using the BEaST patch-based segmentation technique (Eskildsen et al. 2012) which was used to generate skull-stripped brains for subsequent steps. Preprocessed, skull-stripped T1-weighted images were then registered using antsMulitvariateTempalteConstruction tools (<https://github.com/CoBrALab/optimized_antsMultivariateTemplateConstruction>). Briefly, this was used to affinely and nonlinearly register brains to create a consensus average. The initial target was the MNI ICBM NLIN 09c model (1mm isotropic resolution), but a subsequent study-specific average was created, and then upsampled to 0.5mm isotropic. The registration procedure yielded deformation fields from which we can extract absolute Jacobian determinants that encode the contributions from the affine and nonlinear transforms between subject and average, as well as the relative Jacobians which have the affine (linear) component removed. Finally, Jacobians were smoothed with a 2-mm Full Width Half Maximum (FWHM) 3D Gaussian smoothing kernel.

**1.3. Ensuring there is no bias due to relatedness of twin pairs**

To ensure that the relatedness of some of the human subjects did not impact our estimations of sex-differences in brain anatomy, we recomputed our standardized effect sizes in two ways. First, we randomly removed one of the twin pairs from the sample and recomputed the effect size for sex by running the same linear model as described in the main text (see **2.4.1**), which included main effects of sex, mean-centered age, TTV, and Euler number (β_4_). Second, we ran a linear mixed-effects model on the full sample testing again for effects of sex, mean-centered age, TTV, and Euler number, and accounted for relatedness by modeling a family ID as the random intercept. From both of these models, we extracted the standardized effect size (beta-coefficient) for the main effect of sex, and correlated those the effect size for the main effect of sex generated from our main model (**2.4.1**).

**1.4 Sex differences in variance of global and regional brain volume**

We tested for sex differences in the variance of total volume across male and female humans and mice using a Levene’s test. We did so for regional volume as well by residualizing z-scored volumes for TTV, age, and Euler number in humans, and TTV, age, and background strain in mice independently for each sex. We then applied a Levene’s test to test for sex differences in variance across all regions, and corrected for multiple comparisons using FDR correction.

**2. Results**

**2.1 Cross-species sex differences in total grey and total white matter volume**

In addition to sex differences in TTV, we tested for sex differences in total grey matter and total white matter volume across species to understand whether one tissue type was more implicated in driving the sex difference in volume. In humans, males had significantly larger total gray matter volume (t=2.952, p=0.003) while females had significantly larger white matter volume (t=-3.106, p=0.0019) after accounting for TTV, age, and Euler number (***Supplementary Figure 1AC***). In contrast, in mice we observed the opposite relationships, wherein females had significantly larger gray matter volume (t= -2.383, p=0.018) and males had significantly larger while matter volume (t=2.667, p=0.008) after accounting for differences in TTV, age, and background strain (***Supplementary Figure 1BD***).

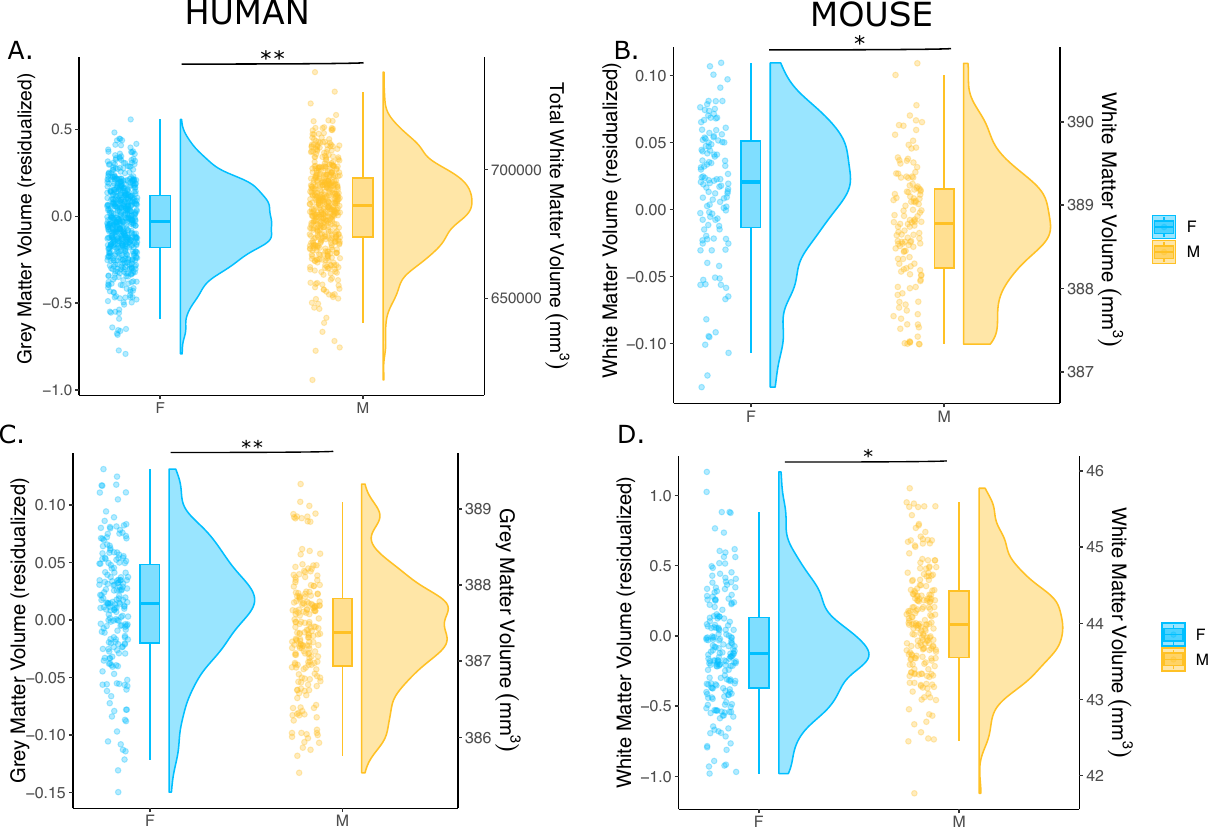
***Supplementary Figure 1. Sex differences in total grey and white matter volume in humans and mice.*** *Sex differences in total grey matter and total white matter volumes are shown for the humans (A: grey; C: white) and mice (B: grey; D: white). Data are represented using individual points, boxplot, and half-violin plot (raincloud plot). **p<0.001; ~p<0.05. M= male, F=female.*

**2.2 Comparison of sex-biased maps when accounting for relatedness in the human sample**

Sex-differences in regional brain volume were not affected by relatedness of twin pairs; both unthresholded and thresholded maps for the effect of sex are identical across our three different models (***Supplementary Figure 1AB***). The random exclusion of one twin pair yielded regional effect size estimates for sex that were highly correlated to those generated from the model that did not exclude a twin pair (r=0.955, t = 228.25, df = 489, p-value < 2.2e-16). Similarly, regional effect size estimates for sex generated from the linear mixed-effects model that accounted for relatedness were highly correlated to those generated from the linear model that did not account for relatedness (r=0.993, t = 190.35, df = 490, p-value < 2.2e-16) (***Supplementary Figure 1CD***).

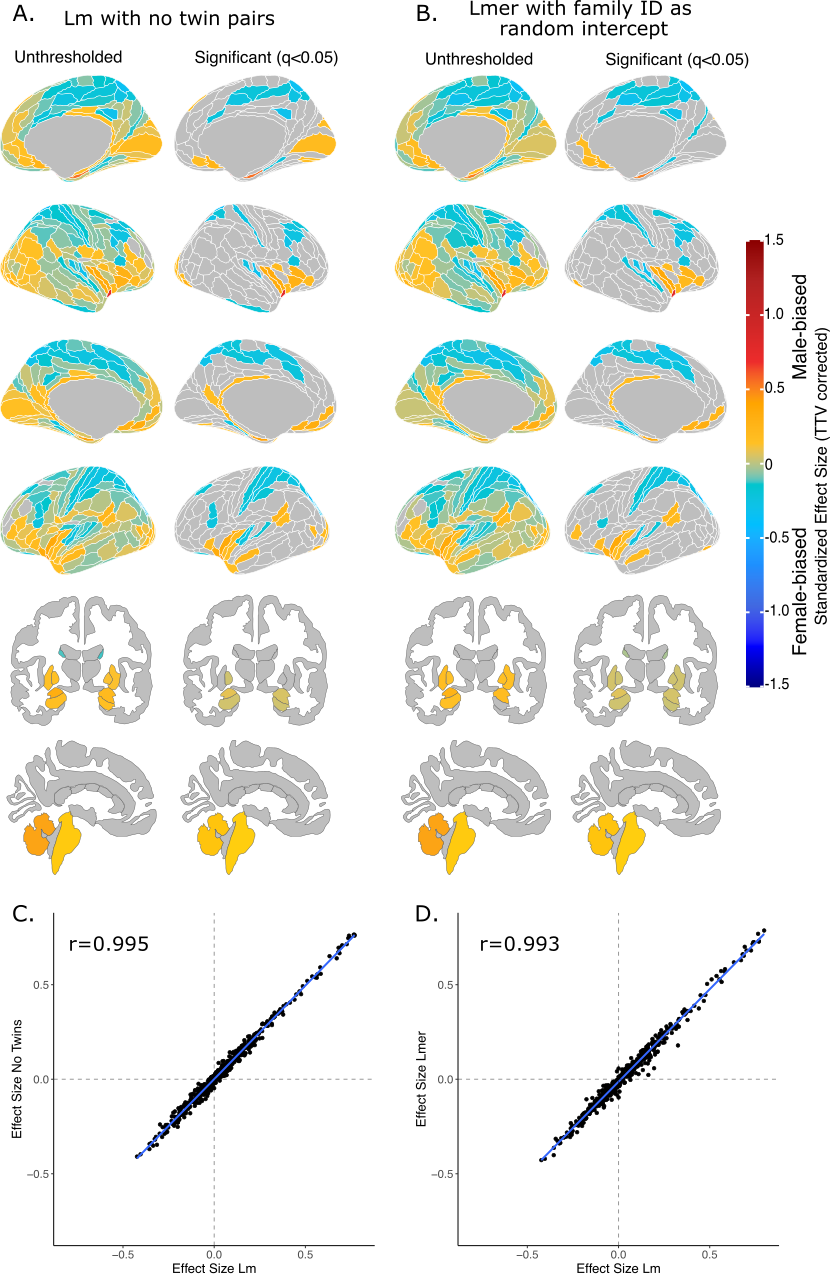

***Supplementary Figure 2. Effect of sex on regional brain volume in humans when accounting for relatedness.*** *Distribution of sex-biased standardized effect sizes for humans when one twin-pair is excluded (A) or when relatedness is accounted for using linear mixed-effects modeling (B). Unthresholded (left) and significant (q<0.05; right) standardized effect sizes for the effect of sex displayed on the human brains. Regions in yellow-red are male-biased and regions in blue are female-biased. (****C****) Correlation of standardized effect sizes for the linear model with no twin pairs compared to that with twin pairs. (****C****) Correlation of standardized effect sizes for the linear model with twin pairs compared to the linear mixed-effects model with twin pairs, but accounting for relatedness.*

**2.3 Sex-biases in regional brain volume in humans and mice without correcting for differences in total tissue volume**

In humans, the male brain is ~13% larger than the female brain, which is reflected in the region-based analyses. There were no female-biased regions, while the mean effect size for the male-biased regions was 0.633 +/- 0.188 (range of 0.200 to 1.283). In contrast, since there were no differences in total brain size between male and female mice, sex differences in regional volumes without correcting for TTV were very similar to those that did correct for TTV (t=2.34, q<0.05). The mean effect size for male-biased regions was still larger than that of the female-biased regions, with a wider range of effect sizes *(Males: median effect size = 0.215, SD=+/-0.263, range=0.006 to 1.073; Females: median effect size = -0.126, SD=+/-0.145, range= -0.0005 to -0.478) (****Supplementary Figure 3****)*.

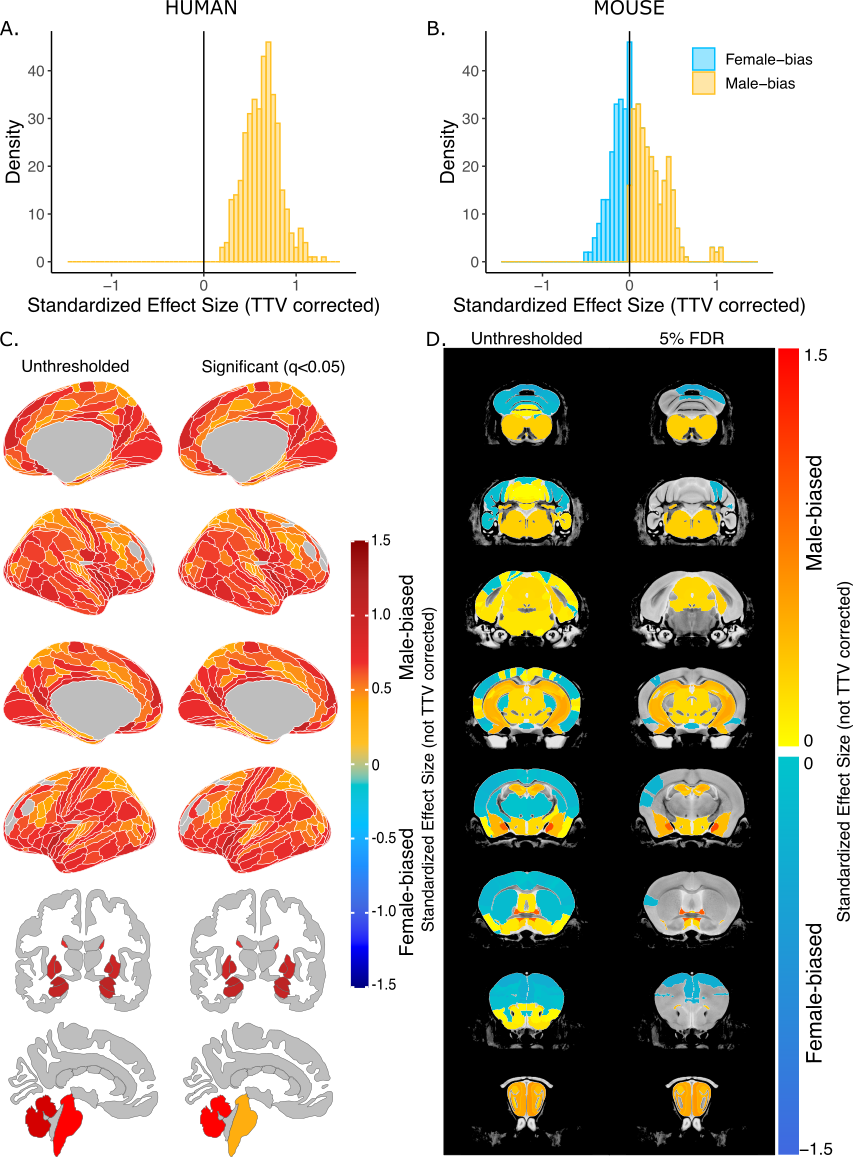

***Supplementary Figure 3. Effect of sex on regional brain volume in humans and mice without TTV correction.*** *Distribution of sex-biased standardized effect sizes for humans (A) and mice (B). Unthresholded (left) and significant (q<0.05; right) standardized effect sizes for the effect of sex displayed on the human (C) and mouse (D) brains. Regions in yellow-red are male-biased and regions in blue are female-biased.*

**2.4 Human males have greater variance in brain volume than females, while mice show no sex differences in variance**

We tested whether there were significant sex differences in the variance of regional volumes in each species and found that in humans, several cortical regions, including the posterior parietal, temporal, frontal opercular, medial prefrontal, posterior cingulate, as well as some subcortical regions (amygdala and hypothalamus) were significantly more variable in human males (q<0.05). In contrast, in mice, there were no regions that had statistically significant sex differences in variance following multiple comparisons correction (q=0.05), however, a handful of regions showed effects at a nominal threshold of p=0.05. Females showed greater variability in the volume of the somatosensory, entorhinal, and visual cortex, as well as hippocampal CA1, while males showed greater variability in some regions of the cerebellum, including the culmen, lingula, and fastigial nuclei, as well as in the olfactory bulbs (***Supplementary Figure 4***). Analyses were repeated without residualizing out the effects of TTV, and in both humans and mice, the results were very similar to the previous analysis (***Supplementary Figure 5***).

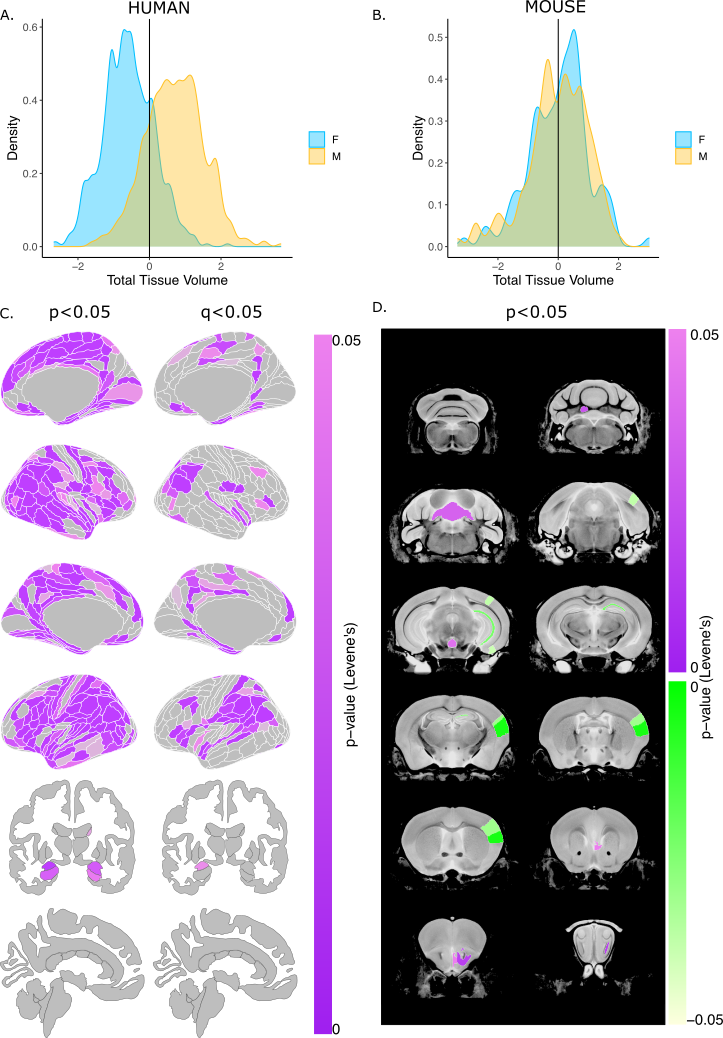

***Supplementary Figure 4. Sex differences in the variability of regional brain volumes (accounting for TTV differences) in humans and mice.*** *Distribution of z-scored total brain volume measures across all humans (A) and mouse subjects (B). Uncorrected (p<0.05; left) and significant (q<0.05; right) sex differences in variability (based on Levene’s test) shown on the human brain Uncorrected (p<0.05) sex differences in variability in the mouse brain (purple = more variable in males; green=more variable in females). Note: all regional human volumes were residualized for TTV, age, and Euler, while regional mouse volumes were residualized for TTV, age, and background strain.*

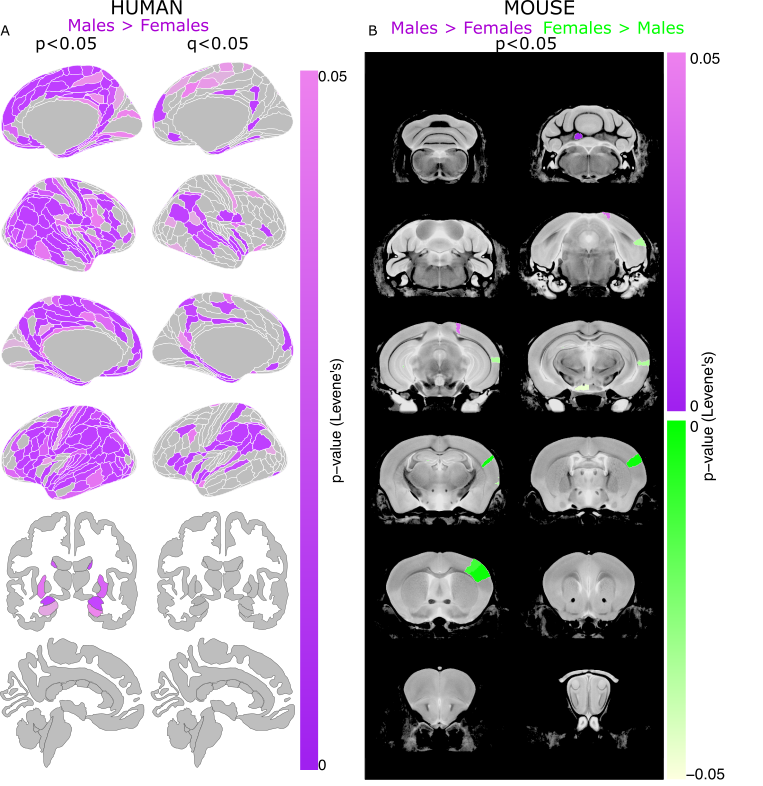

***Supplementary Figure 5***. ***Sex differences in the variability of regional brain volumes (not accounting for TTV differences) in humans and mice.*** *Uncorrected (p<0.05; left) and significant (q<0.05; right) sex differences in variability (based on Levene’s test) shown on the human brain (A) Uncorrected (p<0.05) sex differences in variability in the mouse brain (purple = more variable in males; green=more variable in females) (B). Note: all regional human volumes were residualized for age and Euler, while regional mouse volumes were residualized for age and background strain.*

**2.5 Comparing patterns of sex-biased brain anatomy across species**

Since all effect sizes for humans were male-biased, the cross-species correlation for the effect of sex was not as strong in the analysis that did not covary for TTV, r=0.15 (p=0.25), compared to the TTV-covaried analysis (r=0.30) ***(Supplementary Figure 6A)***. Across cortical regions, there was a low correlation of *r=-0.04* (p=0.86), while for non-cortex the correlation was slightly stronger, but negative, *r=-0.20* (p=0.25). In humans, the subset of homologous regions was all male-biased due to the larger overall brain size in males. In mice, we observed the same patterns of sex-bias as we did in the analyses which contrived for TTV except for the right nucleus accumbens showing no sex bias and a male-bias in the pons (both female-biased in the TTV controlled analysis). The effects are further described in ***Supplementary Table 4.***

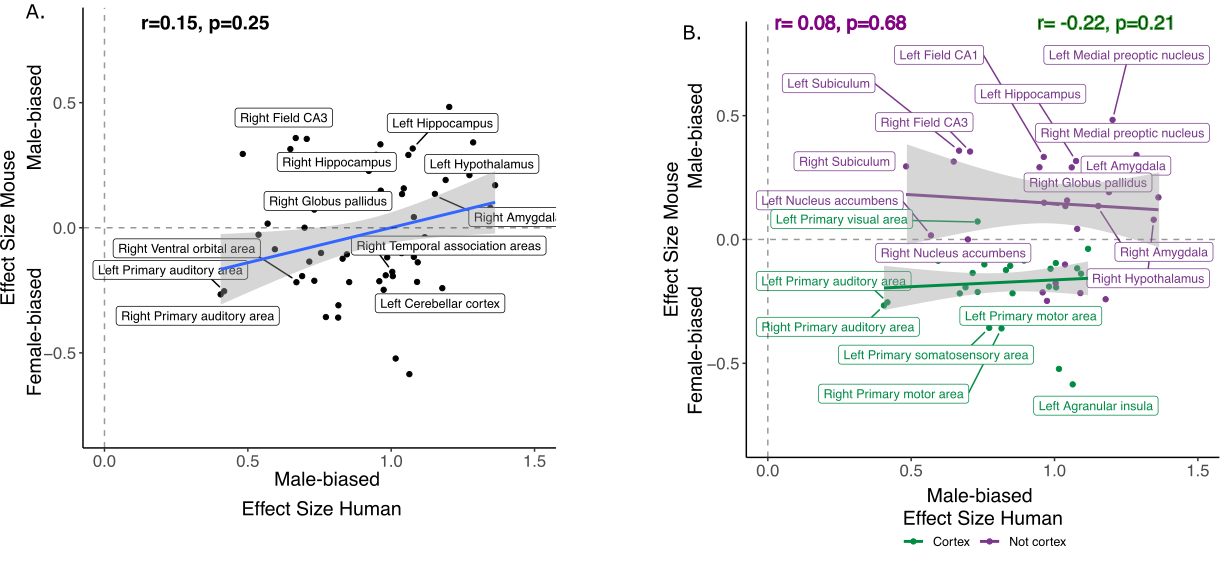

**Supplementary Figure 6. *Correlation of effects of sex in human and mouse homologous brain regions without TTV correction. A****. Standardized effect size correlation for the effect of sex in humans (x-axis) and mice (y-axis) (r=0.09).* ***B****. Correlation of standardized effect sizes for the effect of sex across species for cortical regions (green, r=-0.09), and non-cortical regions (purple, r=0.20).*

***Supplementary Table 4. Mapping of homologous human-mouse brain regions (not TTV-corrected standardized effect sizes).*** *Blue shade highlights female-biased regions, while yellow shade highlights male-biased regions.*

| **Label** | **Glasser/Freesurfer# names** | **Mouse atlas** | **Hemisphere** | **Human Effect Size** | **Mouse Effect Size** |
| --- | --- | --- | --- | --- | --- |
| **Agranular insula** | AVI, AAIC, MI | Agranular insular area | L | **1.025 *** | **-0.484 *** |
|  |  |  | R | **0.985 *** | **-0.453 *** |
| **Amygdala** | Amygdala# | Cortical subplate | L | **1.194 *** | **0.265 *** |
|  |  |  | R | **1.135 *** | **0.240 *** |
| **Anterior cingulate area** | A24pr, a24, p24pr, p24, 24dd, 24dv, p32pr, d32, a32pr, p32, s32 | Anterior cingulate area | L | **0.946 *** | -0.107 |
|  |  |  | R | **0.958 *** | -0.173 |
| **Bed nucleus of stria terminalis** | Bed nucleus of stria terminalis | Bed nucleus of stria terminalis | L | **1.160 *** | **0.937 *** |
|  |  |  | R | **1.131 *** | **0.959 *** |
| **Caudoputamen** | Caudate#, Putamen# | Caudoputamen | L | **0.986 *** | -0.158 |
|  |  |  | R | **0.993 *** | -0.132 |
| **Cerebellar cortex** | Cerebellar cortex# | Cerebellar cortex | L | **1.123 *** | **-0.220 *** |
|  |  |  | R | **1.192 *** | -0.218 |
| **Dentate gyrus, molecular layer** | Dentate gyrus, molecular layer | Dentate gyrus, molecular layer | L | **0.885 *** | **0.292 *** |
|  |  |  | R | **0.935 *** | **0.275 *** |
| **CA1** | CA1 | CA1 | L | **0.936 *** | **0.427 *** |
|  |  |  | R | **0.925 *** | **0.417 *** |
| **CA3** | CA3 | CA3 | L | **0.593 *** | **0.382 *** |
|  |  |  | R | **0.663 *** | **0.460 *** |
| **Entorhinal cortex** | EC | Entorhinal area | L | **0.925 *** | 0.038 |
|  |  |  | R | **0.984 *** | -0.047 |
| **Globus pallidus** | Globus Pallidus# | Pallidum | L | **1.040 *** | **0.167 *** |
|  |  |  | R | **1.049 *** | **0.229 *** |
| **Hippocampus** | Hippocampus# | Hippocampal region | L | **1.045 *** | **0.414 *** |
|  |  |  | R | **1.033 *** | **0.426 *** |
| **Hypothalamus** | Hypothalamus | Hypothalamus | L | **1.374 *** | **0.245 *** |
|  |  |  | R | **1.354 *** | 0.170 |
| **Medial amygdalar nucleus** | Medial amygdalar nucleus | Medial amygdalar nucleus | L | **0.547 *** | **0.926 *** |
|  |  |  | R | **0.654 *** | **1.057 *** |
| **Medial preoptic area** | Medial preoptic area | Medial preoptic area | L | **1.205 *** | **0.472 *** |
|  |  |  | R | **1.275 *** | **0.388 *** |
| **Nucleus accumbens** | Nucleus accumbens# | Striatum ventral region | L | **0.596 *** | 0.079 |
|  |  |  | R | **0.709 *** | 0.106 |
| **Perirhinal area** | PeEc, TF, PHA2, PHA3 | Perirhinal area | L | **0.839 *** | -0.014 |
|  |  |  | R | **0.821 *** | -0.029 |
| **Piriform cortex** | Pir | Piriform cortex | L | **0.988 *** | -0.020 |
|  |  |  | R | **1.076 *** | -0.039 |
| **Posterior parietal association areas** | 5m, 5mv, 5L | Posterior parietal association areas | L | **0.519 *** | 0.028 |
|  |  |  | R | **0.554 *** | 0.066 |
| **Primary auditory area** | A1 | Primary auditory area | L | **0.352 *** | -0.192 |
|  |  |  | R | **0.357 *** | -0.183 |
| **Primary motor area** | 4 | Primary motor area | L | **0.785 *** | **-0.237 *** |
|  |  |  | R | **0.795 *** | **-0.269 *** |
| **Primary somatosensory area** | 1, 2, 3a, 3b | Primary somatosensory area | L | **0.743 *** | **-0.332 *** |
|  |  |  | R | **0.671 *** | -0.162 |
| **Primary visual area** | V1 | Primary visual area | L | **0.729 *** | 0.098 |
|  |  |  | R | **0.703 *** | -0.085 |
| **Retrosplenial area** | RSC | Retrosplenial area | L | **0.838 *** | 0.010 |
|  |  |  | R | **0.770 *** | 0.062 |
| **Subiculum** | PreS | Subiculum | L | **0.643 *** | **0.370 *** |
|  |  |  | R | **0.433 *** | **0.382 *** |
| **Temporal association areas** | FFC, PIT, TE1a, TE1p, TE2a, TF, STV, STSvp, STSva | Temporal association areas | L | **1.084 *** | 0.159 |
|  |  |  | R | **1.060 *** | 0.024 |
| **Thalamus** | Thalamus# | Thalamus | L | **1.019 *** | -0.040 |
|  |  |  | R | **0.994 *** | -0.087 |
| **Ventral orbital area** | 10r, 10v | Ventral orbital area | L | **0.666 *** | -0.121 |
|  |  |  | R | **0.605 *** | -0.093 |
| **Brain stem (midline)** | Brainstem# | Midbrain, Hindbrain | M | **1.209 *** | **0.226 *** |
| **Medulla (midline)** | Medulla | Medulla | M | **1.111 *** | **0.255 *** |
| **Midbrain (midline)** | Midbrain | Midbrain | M | **1.280 *** | **0.194 *** |
| **Pons (midline)** | Pons | Pons | M | **1.101 *** | 0.101 |

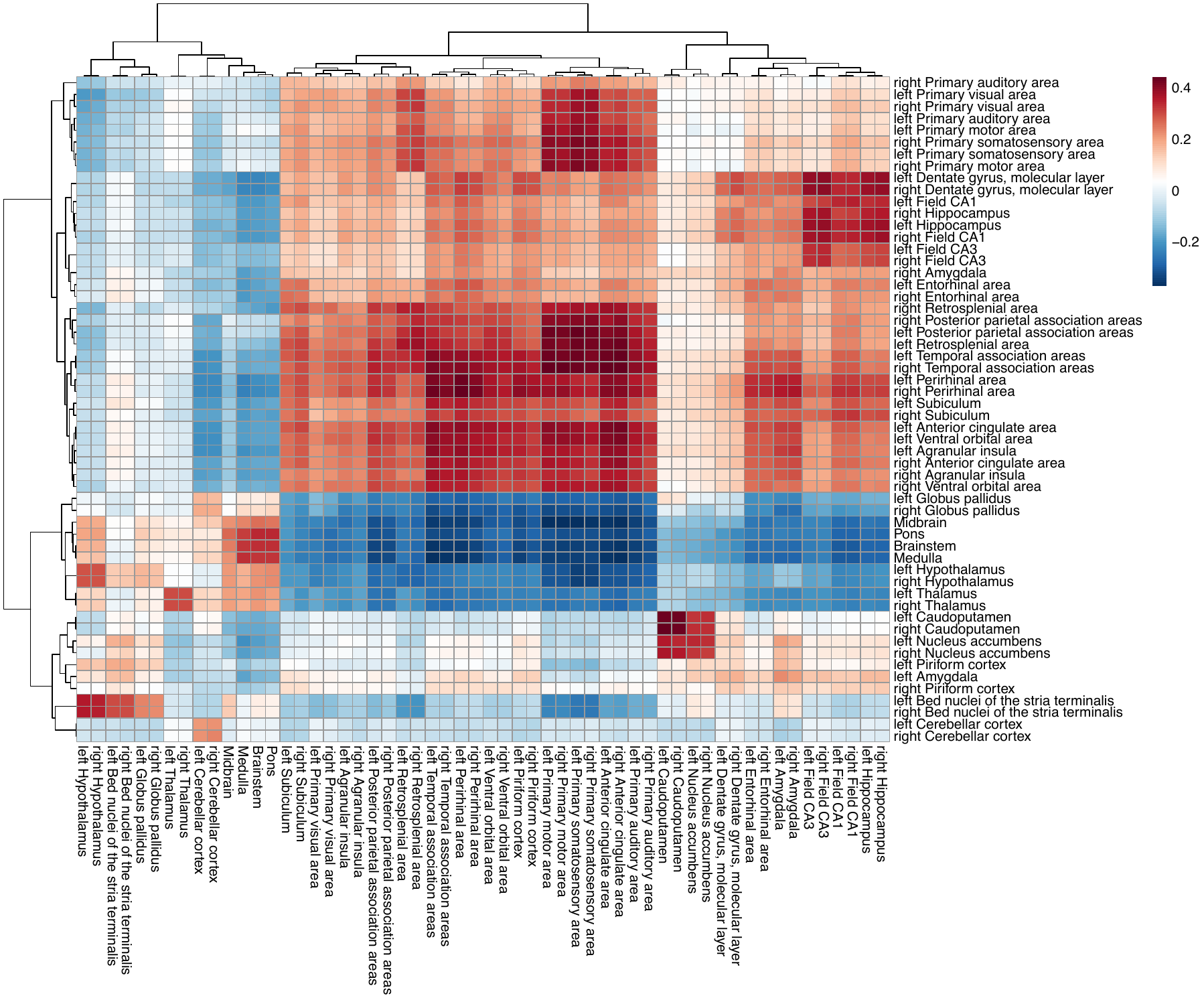

***Supplementary Figure 7. Similarity matrix for cross-species homologous gene expression.*** The matrix displays the correlation between 56 homologous human (x-axis) and mouse (y-axis) brain regions based on the expression of 2835 homologous genes. Hierarchical clustering was used to order the rows and columns using the R function *pheatmap.*

**2.6 Correlating the anatomical sex effect similarity score to the similarity of homologous gene expression across homologous brain regions**

We limited our homologous genes to those that were only expressed on the X-chromosome to determine whether their regional expression would be more highly correlated to patterns of sex-biased neuroanatomy. The robust correlation between anatomical sex similarity score and transcriptional similarity of X-chromosome genes (n=91) was slightly stronger, although very similar to that of the full homologous gene set (n=2835), *r=0.25* (p=0.07). As with the full gene set, the relationship was stronger, and statistically significant, for cortical regions, *r=0.62* (p=0.0007), than non-cortical regions, *r=0.30* (p=0.11).

Filtering genes associated with sex steroid hormones-based Gene Ontology annotations (n=30) yielded a weak, and negative correlation, *r=-0.11* (p=0.44). Interestingly, the relationship for cortical regions was positive and similar to the relationships described above, *r=0.29*, (p=0.14), while the correlation in the subcortex was negative, *r=-0.13* (p=0.50) (***Supplementary Table 7*** for gene lists with GO terms and ***Supplementary Table 8*** for those lists filtered to include only homologous genes*)*. Lastly, we examined whether limiting sex hormone genes to more male-biased (androgen signaling genes, n=11) or more female-biased (estrogen and progesterone signaling genes, n=23) genes affected the correlation between anatomical sex similarity score and gene expression similarity. We found that androgen genes were positively correlated (r=0.23, p=0.08) with the anatomical sex similarity score, with a weak correlation for cortex (r=0.05, p=0.81) but a strong one for non-cortex (r=0.46, p=0.01). In contrast, estrogen and progesterone genes were negatively correlated with the anatomical sex similarity score (r=-0.21, p=0.13), with a positive correlation in the cortex (r=0.29, p=0.15) and a negative relationship in the non-cortex (r=-0.27, p=0.15).

Finally, we recomputed the transcriptional similarity by randomly resampling various subsets of homologous genes 10,000 times, and then correlated those similarity values to the anatomical sex congruence across regions. The subsets were chosen to reflect the number of genes selected for the various hypothesis-driven subsets, i.e., n=91 for X-chromosome genes, n=34 for sex hormone genes, n=11 for androgen genes, and n=23 for estrogen and progesterone genes. For the distribution based on 91 genes we observed a mean r of 0.05 (min=-0.20, max=0.32); for 34 genes we observed a mean r of 0.04 (min=-0.34, max=0.39); for 11 genes we observed a mean r of 0.03 (min=-0.42, max=0.46), and for 23 genes we observed a mean r of 0.04 (min=-0.33, max=0.38). Relative to the null distribution of correlated values, the relationship anatomical sex congruence and transcriptional similarity were significant for the subset of X-chromosome genes (n=91, p_null_=0.0014), trending towards significant for the subset of sex hormone genes (n=34, p_null_=0.08),and for androgen genes (n=11, p_null_=0.06), and significant for the subset of estrogen and progesterone genes (n=23, p_null_=0.01).

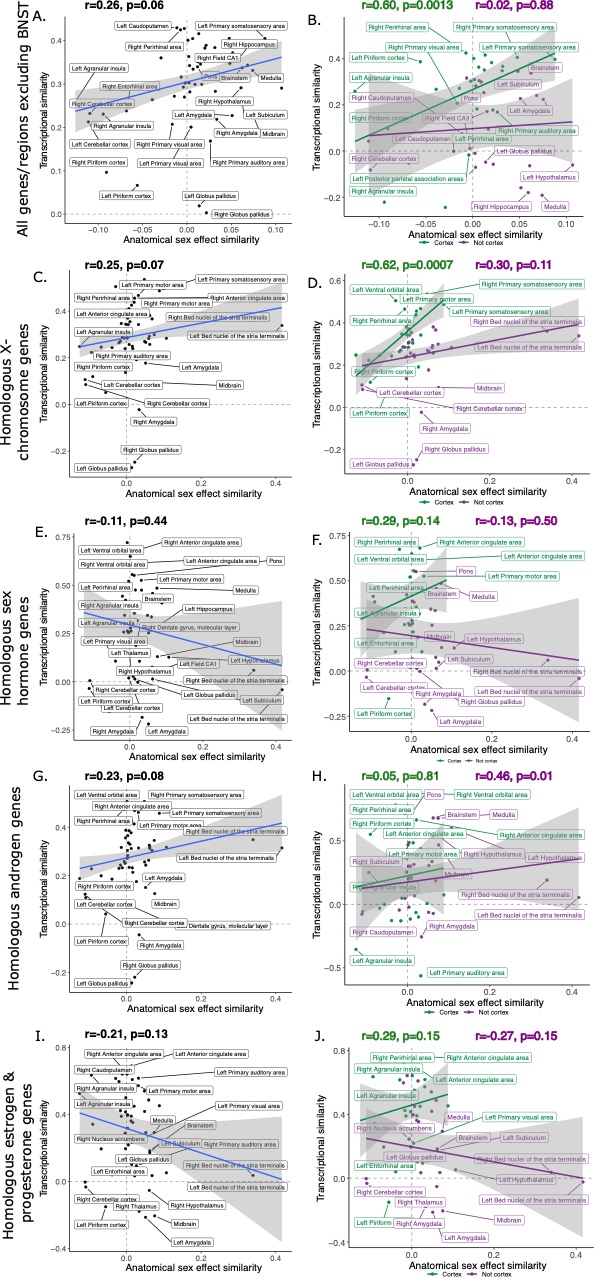

***Supplementary Figure 8. Robust correlation of anatomical similarity of sex effects and transcriptional similarity of homologous brain regions across species.*** *Correlation of anatomical similarity and transcriptional similarity using all homologous genes but excluding the BNST (****AB****), only X-chromosome homologous genes (n=91;* ***CD****), using only sex hormone genes (androgen, estrogen, progesterone genes, n=30;* ***EF****), just androgen genes (****GH****), or just estrogen and progesterone genes (****IJ****) across all homologous regions or split into cortical and non-cortical.*

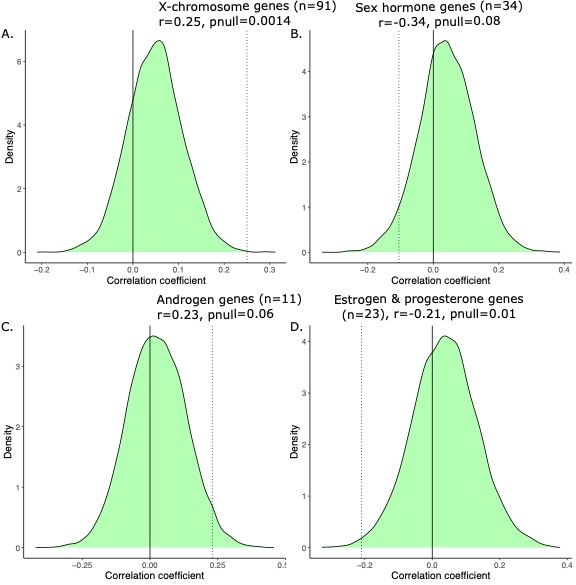

***Supplementary Figure 9. Generating a null distribution of correlation coefficients for the anatomical vs transcriptional similarity.*** *Correlations are significant relative to null distribution (shown in green) generated by randomly sampling subsets of 2835 homologous genes corresponding to the biologically informed subsets, recomputing the transcriptional similarity, and correlating that with the anatomical similarity 10,000 times. We have subsets of 91 genes in (****A****) corresponding to X-linked genes, 34 genes in (****B****) corresponding to sex hormone genes, 11 genes in (****C****) corresponding to androgen games, and 23 genes in (****D****) corresponding to estrogen and progesterone genes. For each, the observed correlation was compared to the null correlation to generate a p-value, displayed on the graph.*
